# ZapLine: a simple and effective method to remove power line artifacts

**DOI:** 10.1101/782029

**Authors:** Alain de Cheveigné

## Abstract

Power line artifacts are the bane of animal and human electrophysiology. A number of methods are available to help attenuate or eliminate them, but each has its own set of drawbacks. In this brief note I present a simple method that combines the advantages of spectral and spatial filtering, while minimizing their downsides. This method is applicable to multichannel data such as electroencephalography (EEG), magnetoencephalography (MEG), or multichannel local field potentials (LFP). I briefly review past methods, pointing out their drawbacks, describe the new method, and evaluate the outcome using synthetic and real data.

## Introduction

Most laboratories are criss-crossed with power lines, and peppered with line-powered lights and electronic equipment. The electric and magnetic fields that these create can interfere with the tiny potentials and fields that are measured from the brain with techniques such as electroencephalography (EEG), magnetoencephalography (MEG) or other electrophysiological techniques. Power line artifacts are manifest in the spectrum as a series of narrow peaks at multiples of the power line frequency (50 Hz or 60 Hz depending on the geographical region) (Fig. 1). The mechanisms by which that power makes its way into the rig can be puzzling for the experimentalist (ground loops, capacitive coupling, etc.), and the presence of some amount of line artifact in the data is often accepted as a fact of life. Line interference is usually unavoidable in MEG, which is sensitive to fields from distant sources, and a scientist often comes across preexisting datasets for which the question of ‘avoiding artifacts at the source’ is moot.

**Figure 1:**
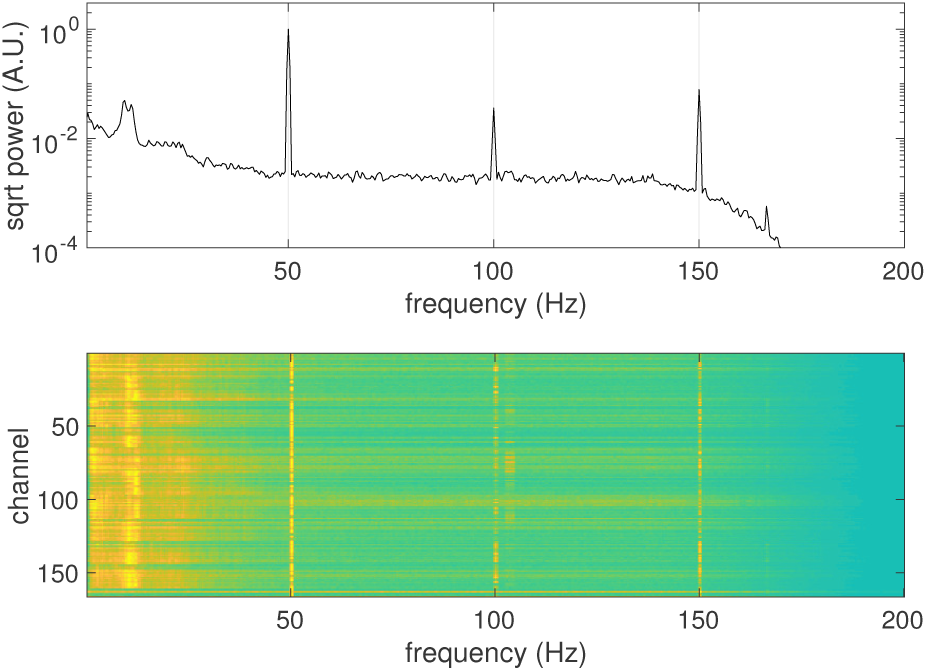
Square root power spectral density (PSD) of a MEG data file. Power line artifacts are manifest as spectral peaks at the power line frequency (here 50 Hz) and multiples. The bottom panel shows the same information per channel as a colour plot. All channels are contaminated to some degree.

A wide range of approaches have been proposed to attenuate or remove line artifacts, including *lowpass* and *notch filters* (Luck, 2005), *frequency domain filters* (Mitra and Pesaran, 1999; Mullen, 2012; Keshtkaran and Yang, 2014; Leske and Dalal, 2019), *regression based on a reference signal* (Vrba and Robinson, 2001; de Cheveigné and Simon, 2007), *independent component analysis (ICA)* (Barbati et al., 2004; Escudero et al., 2007) or other *spatial filtering* techniques (de Cheveigné and Parra, 2014). These methods are well known and widely used, and there are ongoing efforts to develop new ones (Leske and Dalal, 2019). Many of these methods are listed in standard guidelines and textbooks (Picton et al., 2000; Widmann and Schröger, 2012; Luck, 2005; Gross et al., 2013), implemented in widely-used data analysis toolboxes (Delorme and Makeig, 2004; Oostenveld et al., 2011; Gramfort et al., 2014), or integrated in automated processing pipelines (Bigdely-Shamlo et al., 2015; Gabard-Durnam et al., 2018).

A straightforward solution is simply to low-pass filter the data with a cutoff below the line frequency (50 or 60 Hz). The primary motivation is often to enhance slowly varying features relative to fast fluctuations judged less relevant, line artifact reduction being an added benefit. A cutoff in the range 10 - 30 Hz is typical (Fig. 2 top). Downsides are (a) loss of high frequency information, (b) limited attenuation at line frequency unless the cutoff is steep, and (c) waveform distortion, particularly if the cutoff is steep and the filter impulse response consequently long (Widmann and Schröger, 2012; Cohen, 2014; de Cheveigné and Nelken, 2019).

**Figure 2:**
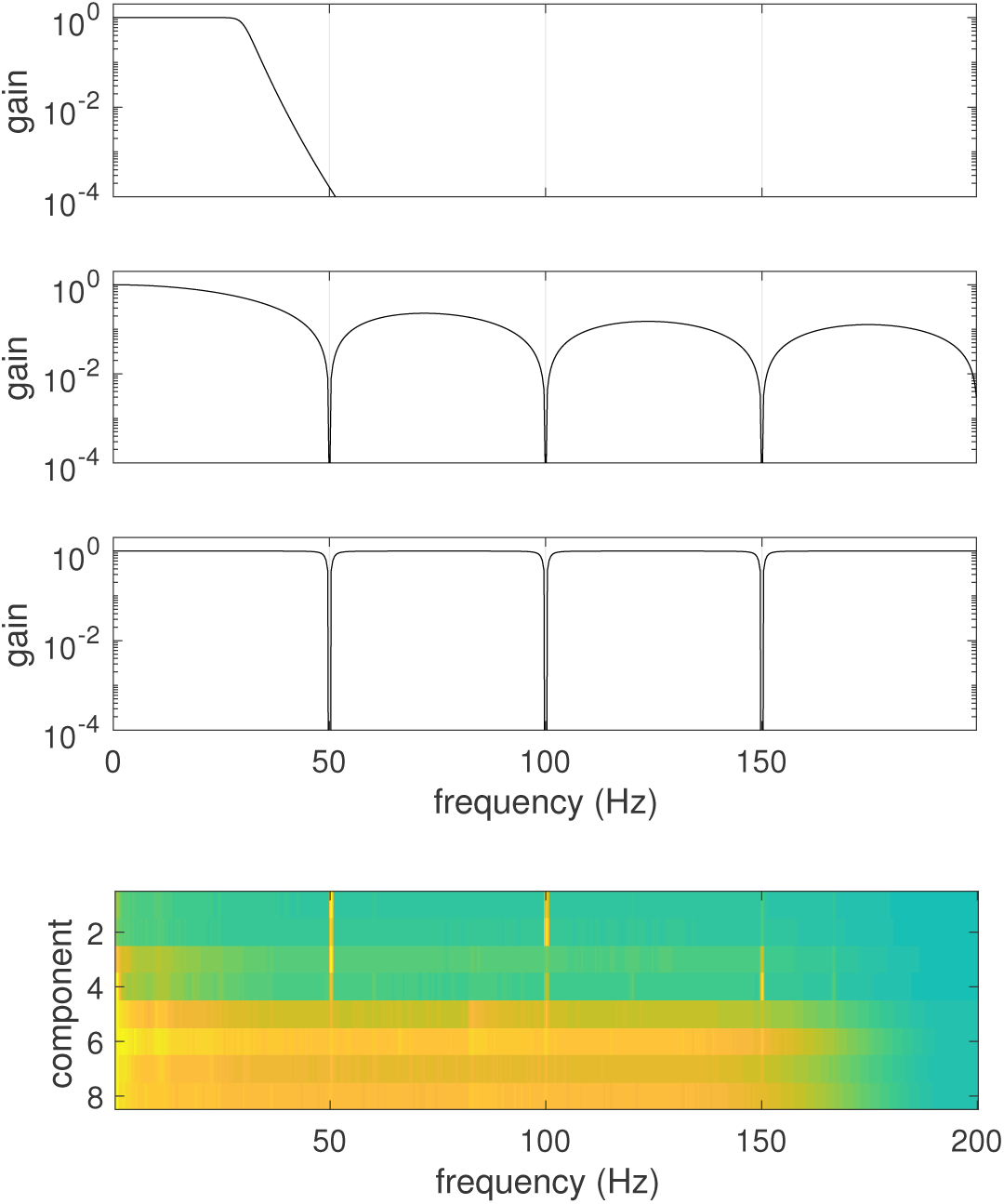
Top panel: magnitude transfer function of a lowpass filter (Butterworth, order 16, cutoff 30 Hz). Second panel: transfer function of a smoothing filter with square kernel of duration 1/50 Hz. Third panel: transfer function of a multiple notch filter with notches at 50 Hz and multiples. Each notch is implemented as a 2nd order IIR filter with 0.01 relative bandwidth. Bottom panel: PSDs of the first 8 components of a DSS analysis of a 166-channel MEG data file designed to find dimensions dominated by power line noise. The first 4 components, clearly dominated by power line noise, can be projected out of the data to obtain clean data.

One effective version of this idea convolves the data with a square-shaped impulse response with a duration exactly equal to the line period. The transfer function of this filter has zeros at the line frequency and all harmonics, and thus the filter *perfectly* removes the artifact (Fig. 2 panel 2), and the impulse response is short so waveform smearing or ringing are minimized. In the event that the sampling rate is not multiple of the line frequency, a reasonably effective ‘square’ impulse response can be approximated by interpolation (de Cheveigné and Arzounian, 2018).

Alternatively, a notch filter, or set of notch filters, can be applied to attenuate the signal at the line frequency and harmonics. Attenuation at other frequencies is minimal if the notches are narrow (Fig. 2 panel 3). The downside is that the very long impulse response of a notch filter can lead to artifacts triggered by the onset, offset, or transitions within the data (Vale-Cardoso and Guimarães, 2009; Kiraç et al., 2015). Analogous to applying a notch filter is to fit to the waveform a quadrature pair of sinusoids at the line frequency, and subtracting the fit. Additional pairs can be fit to all harmonics. The same result is obtained in the frequency domain by zeroing the corresponding components, as long as the sampling rate is a multiple of the line frequency, but the fit-and-subtract approach is more flexible in that it allows arbitrary frequency ratios, and the fit can be weighted to avoid contamination by glitches (de Cheveigné and Arzounian, 2018). Regardless of the method, fitting a sinusoid is ineffective if the power line components fluctuate in amplitude and/or phase.

An alternative to time-domain or frequency-domain filtering is to project the data on a reference channel that picks up only line components, and subtract the projection. Certain MEG systems are equipped with reference channels to support this approach, which also allows suppressing a wider range of artifacts (e.g. slow fields due to machinery) (Vrba and Robinson, 2001; de Cheveigné and Simon, 2007). If no reference channel is available, one can be ‘created’ from multichannel data by applying a spatial filter designed to isolate line-dominated components. ICA has been proposed for this purpose (Barbati et al., 2004; Escudero et al., 2007), and other data-driven techniques are also effective, for example the JD (Joint Diagonalization) method also referred to as DSS, Denoising Source Separation (de Cheveigné and Parra, 2014). Figure 2 (bottom) shows, as a colour plot, the PSD of the first few components of a JD/DSS analysis designed for this purpose. The first 4 of these, strongly dominated by line components, may be used as ‘reference channels’ to project out line artifacts (de Cheveigné and Parra, 2014).

The drawback of the spatial filtering approach is that the power line components are not always linearly separable from brain activity, so projecting out the line-dominated components may lead to loss of useful data. It may also happen that those components are contaminated by other artifacts (such as glitches or drifts), that are then injected into previously uncontaminated channels.

The algorithm described in this paper addresses these issues, combining the benefits of both spatial and temporal filtering.

## Methods

### Data model

Data consist of a time series matrix **X** of dimensions *T* (time) *× J* (channels) that is assumed to be the sum

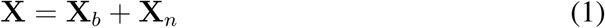

of linear transforms of brain sources **B** and power line noise sources **N**. These sources and transforms are unknown. Linear analysis methods such as ICA try to find a ‘demixing matrix’ to apply to the data to reverse the mixing, but this is only possible if they are linearly separable within the data. Here we assume that this is *not* the case: there exists no scalar demixing matrix **D** such that **X**_*b*_**D** = **X**_*b*_ and **X**_*n*_**D** = 0. Likewise, spectral filtering methods try to find a filter that isolates sources one from another, but here we assume that they are *not* separable in the spectral domain, i.e. there exists no filter *ℱ* such that **X**_*b*_* *ℱ* = **X**_*b*_ and **X**_*n*_* *ℱ* = 0. However, we do assume that there exists a *spatio-spectral* transform in the form of a multichannel filter *𝒯* such that brain and line noise *are* separable, **X**_*b*_ * *𝒯*= **X**_*b*_ and **X**_*n*_ * *𝒯* = 0, at least approximately. Our aim is to find such a transform.

### Algorithm

In a first step, **X** is convolved with a square-shaped smoothing kernel *𝒦* of duration 1*/f*_*line*_ where *f*_*line*_ is the line frequency. This implements a smoothing (low-pass) filter with zeros at *f*_*line*_ and all multiples (as illustrated in Fig. 2 panel 2). Calling **X**^′^= **X** * *𝒦* the filtered signal and **X**″ the remainder, we have

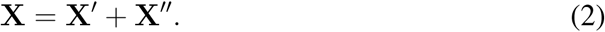

The data are thus decomposed into a line-suppressed and a line-dominated part, the sum of which perfectly reconstitutes the signal.

In a second step, a *spatial* filter **D** is applied to the line-dominated part only, to project out the spatial dimensions that carry line-dominated power. The spatial filter **D** is estimated using the JD/DSS method (de Cheveigné and Parra, 2014), using, as a bias filter, a frequency domain filter with peaks at 1*/f*_*line*_ and multiples. More precisely, the JD/DSS analysis produces a transform matrix **M** such that the first column of **X**′**M** is the linear transform with the highest possible ratio of line power to total power, the second column has the highest ratio within the subspace orthogonal to the first, and so-on. Line power is thus concentrated in the first *d* components. The spatial filter matrix **D** is then formed as the product of the last *J - d* columns of **M** multiplied by the last *J-d* rows of its pseudoinverse. Multiplying **X**^′^ by **D** effectively projects out the *d* dimensions most dominated by line noise.

The clean signal is then reconstructed as

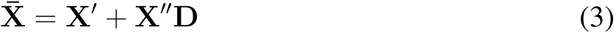

It is easy to see that 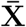 contains *no* power line components, as those components are perfectly cancelled in the first term of Eq. 3 by a spectral filter and in the second term by a spatial filter. Other than that, the data are barely affected: the 1/*f*_*line*_ square-shaped smoothing kernel is applied only to *d* out of *J* spatial components, the others *J - d* remaining unfiltered, while the spatial filter **D** is applied only to the line-dominated, high pass part portion **X***″* of the data, the line-free low pass part remaining untouched. In contrast to spectral filtering methods, the clean data are *full-band*. In contrast to spatial filtering methods, they are *full rank*.

### Implementation

Processing is implemented by the nt_zapline() function of the NoiseTools toolbox (audition.ens.fr/adc/NoiseTools/). Example scripts are available at http://audition.ens.fr/adc/NoiseTools/src/NoiseTools/EXAMPLES/zapline/.

The algorithm has one main parameter: *d*. An additional ‘hidden’ parameter is the bias filter used for JD in the second stage. By default, that filter is implemented in the Fourier domain using an 1024-point FFT by selecting bins at the line frequency and multiples, and setting all others to zero. That filter serves only as a bias: it does not affect the cleaned data.

## Results

### Simulated data

Simulated multichannel data consisted of a 10000 × 100 matrix of normally distributed random samples, taken to represent a wide-band ‘signal’, to which was added a periodic waveform (sinusoid half wave rectified and raised to power 3) multiplied by a 1 × 100 random matrix, taken to represent ‘line noise’. Signal and noise were scaled to approximately the same power. The algorithm was applied with *d* = 1. Figure 3 (top) shows the square root PSD of the mixture (left), and of the denoised data (right, green). The part removed, attributed to line noise, is plotted in red. The square root PSD of the denoised data is flat, as expected for uncorrelated noise, and contains no trace of the simulated line noise. The line noise instead shows prominently in the part removed. The spectral ‘floor’ of the part removed is about one order of magnitude below the denoised signal in square root power, which translates to two orders of magnitude in power. This indicates that the amount by which the signal was changed at frequencies other than line and harmonics is minimal, so any deleterious effect of processing is small.

**Figure 3:**
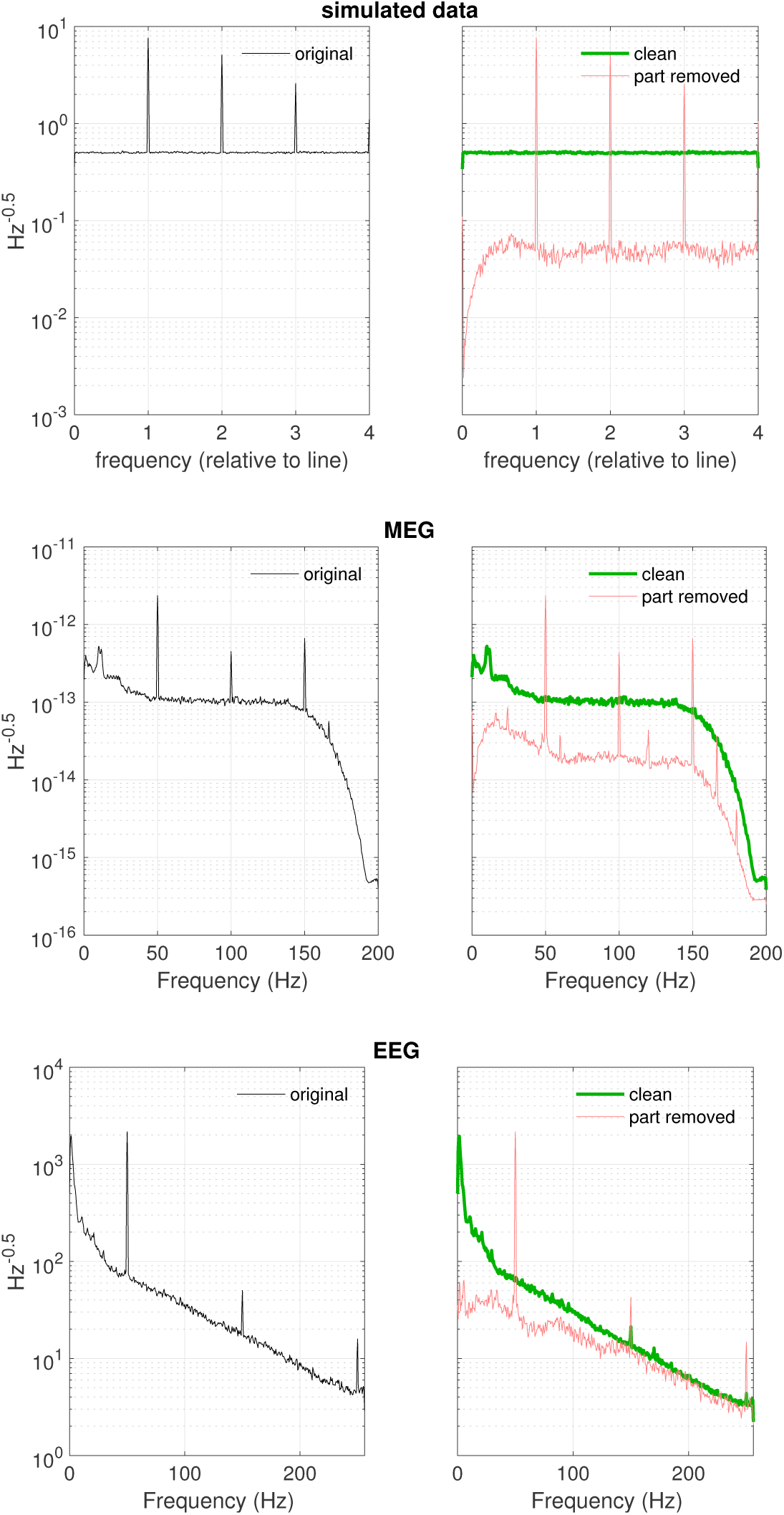
Square root PSD of line-contaminated data (left), cleaned data (right, green), and residual (right, red). Top: simulated data, middle: EEG, bottom: MEG.

### EEG

EEG data consisted of 136 channels of data from a BioSemi system sampled at 512 Hz, taken from a study on auditory attention (Di Liberto et al., 2015). The square root PSD (Fig. 3 middle left) shows a prominent peak at 50 Hz and odd harmonics, that are prominent in the part removed (right, red) but absent in the denoised data (right, green). The algorithm was here applied with *d* = 2 (two spatial dimensions were removed from **X***″*).

### MEG

MEG data consisted of 166 channels of data from a Yokogawa system downsampled to 200 Hz. These data are typical of MEG data in that they are strongly dominated by power line noise (Fig. 3 bottom left). Power line components are absent in the denoised data (right, green) but prominent in the part removed (right, red). Interestingly, the analysis reveals additional weak spectral lines, some of which seem to reflect aliasing, that were not visible in the original data. The algorithm was here applied with *d* = 4 (four spatial dimensions were removed from **X***″*).

Additional examples, together with code and links to data, are provided at http://audition.ens.fr/adc/NoiseTools/src/NoiseTools/EXAMPLES/zapline/. They help make the point that the method is effective and safe for a wide range of data.

## Discussion

The algorithm is both highly effective at removing power line artifacts, and respectful of non-artifactual parts of the signal. It is computationally cheap and easy to apply, with only one main parameter, *d*, that specifies the number of spatial components to reject from the power-line dominated part of the data **X***″*. Multiple power line components indicate multiple sources of artifact, each with a different time course and spatial signature. MEG tends to have more power line components than other techniques, reflecting its sensitivity to distant sources. The appropriate value for *d* is usually 1 or a small number. Choosing *d* too small makes the algorithm less effective because not all contaminated dimensions are removed. Choosing it too large will cause more spatial dimensions to be affected by the line-suppression filter than needed, which is usually of little consequence. The method is thus attractive as a ‘no-nonsense’ line-suppression stage within an automated proceessing pipeline.

Power line artifacts may be detrimental in that they mask true brain activity. They may also defeat or interfere with processing methods designed to reveal that activity. For example, in an evoked-response experiment, if the timing of trials is inadvertently set such that each trial begins at the same phase of the power line cycle, the line artifact adds up in phase instead of washing out. Worse, if an algorithm such as DSS is used to isolate repeatable directions within the data, if may return artifact-dominated (or artifact-contaminated) directions. There is also an indirect effect, in that line artifacts motivate processing steps (filtering, etc.) that may distort brain activity patterns.

Lowpass filtering has the obvious drawback that it suppresses high-frequency patterns, as well as a more insidious effect of distorting brain waveforms (de Cheveigné and Nelken, 2019). Notch filtering has less of a spectral footprint, but the very long impulse response may trigger artifacts at onsets, offsets, and changes in the signal (Vale-Cardoso and Guimarães, 2009; Kiraç et al., 2015). Regression and frequency-domain methods that fit sinusoids to the data are sensitive to glitches and variations of phase and amplitude of the power line signals.

Spatial filtering methods face a different set of issues. ICA exploits non-Gaussianity, non stationarity, or non-whiteness of sources mixed within the data to find a demixing matrix that isolates them. Isolated components then need to be classified according to the kind of source they represent, for example based on their spectral properties, and removed to obtain clean data. However, the selection process is hard to automate, and furthermore line artifacts may not always be perfectly resolved among the components. Spatial filtering based on DSS/JD is easier to deploy and possibly more effective in that the components are directly optimized to maximize power at line frequencies, rather that merely non-gaussianity (de Cheveigné and Parra, 2014). Regardless of which method is used to identify the line-dominated dimensions to remove, the cleaned data are rank-deficient, which may be a problem for certain algorithms (e.g. some versions of ICA) that expect data to be full-rank. Brain activity that happens to fall within those dimensions is lost, and artifacts that occur within those dimensions may be injected into otherwise clean data.

The method proposed here draws on both spatial and spectral filtering. The problems of the former are mitigated by restricting it to a limited part of the spectrum, and those of the latter by limiting it to a small subset of dimensions within the data. If need, the particular smoothing filter *𝒦* described in the Methods could be replaced by some other line-suppressing filter, e.g. low-pass or notch.

The principle behind the algorithm (Fourier-filter into two complementary paths and apply different spatial filtering to each) is possibly of wider applicability, for example to suppress or enhance narrowband activity such as alpha while minimizing effects on other brain components. Processing amounts to application of a particular multichannel FIR (finite impulse response) filter that the algorithm discovers. More general ways to find such filters have been suggested (de Cheveigné, 2010), but their greater flexibility is not necessarily an advantage as it can lead to overfitting. The present method is more tightly constrained to do ‘the right thing’.

There is recent interest in real-time methods to process EEG for applications such as steering a hearing aid. The present method is ideal in that context in that it involves filtering with a FIR filter of very short impulse response (hence low latency), and a spatial filter that can easily be learned on past data, and if necessary updated to accommodate changes in mixing weights of the line artifact.

## Conclusion

I presented a simple, effective, and efficient method to remove power line artifacts from multichannel data, applicable to EEG, MEG and other multichannel electrophysiology techniques. The method combines spectral and spatial filtering in such a way as to attain perfect artifact rejection while minimizing deleterious effects. The method requires only minimal tuning, and is therefore an ideal component to include in an automatized data analysis pipeline.

## Acknowledgements

This work was supported by grants ANR-10-LABX-0087 IEC, ANR-10-IDEX-0001-02 PSL, and ANR-17-EURE-0017.

